# Differential Effects of “Resurrecting” Csp Pseudoproteases during *Clostridioides difficile* Spore Germination

**DOI:** 10.1101/855569

**Authors:** M. Lauren Donnelly, Emily R. Forster, Amy E. Rohlfing, Aimee Shen

**Author notes:** Address correspondence to Aimee Shen. Abbreviations List: n/a.

## Abstract

*Clostridioides difficile* is a spore-forming bacterial pathogen that is the leading cause of hospital-acquired gastroenteritis. *C. difficile* infections begin when its spore form germinates in the vertebrate gut upon sensing bile acids. These germinants induce a proteolytic signaling cascade controlled by three members of the subtilisin-like serine protease family, CspA, CspB, and CspC. Notably, even though CspC and CspA are both pseudoproteases, they are nevertheless required to sense germinants and activate the protease, CspB. Thus, CspC and CspA are part of a growing list of pseudoenzymes that play important roles in regulating cellular processes. However, despite their importance, the structural properties of pseudoenzymes that allow them to function as regulators remain poorly understood. Our recently determined crystal structure of CspC revealed that its degenerate site residues align closely with the catalytic triad of CspB, so in this study we tested whether the ancestral protease activity of the CspC and CspA pseudoproteases could be “resurrected.” Restoring the catalytic triad to these pseudoproteases failed to resurrect their protease activity, although the mutations differentially affected the stability and function of these pseudoproteases. Degenerate site mutations destabilized CspC and impaired spore germination without impacting CspA stability or function. Thus, our results surprisingly reveal that the presence of a catalytic triad does not necessarily predict protease activity. Since close homologs of *C. difficile* CspA occasionally carry an intact catalytic triad, our results imply that bioinformatics predictions of enzyme activity may overlook pseudoenzymes in some cases.

## Introduction

Catalytically-deficient structural homologs of functional enzymes known as pseudoenzymes were first discovered more than 50 years ago, but they were frequently dismissed as vestigial remnants of evolution because they lacked catalytic activity [1]Ribeiro, 2019 #94}. However, the prevalence of pseudoenzyme genes, estimated at 10-15% of a typical genome [2] across all domains of life [1, 3], suggested that they have important biological functions. Indeed, recent work has established that pseudoenzymes perform diverse and crucial cellular functions [1, 4, 5], controlling metabolic and signaling pathways in processes ranging from cell cycle progression to protein trafficking. In this sense pseudoenzymes have been ‘brought back to life’[1, 3-5], which is why they have been referenced as ‘zombie’ proteins in the literature [1, 4].

While pseudoenzymes have been identified in over 20 different protein families, including pseudokinases, pseudophosphatases, and pseudoproteases [1, 3], the mechanisms by which they modulate cellular processes are poorly understood, since relatively few predicted pseudoenzymes have been thoroughly studied in biological systems. Studies thus far indicate that pseudoenzymes can allosterically regulate the activity of cognate enzymes, nucleate protein complexes by acting as cellular scaffolds, control protein localization, and act as competitors for substrate binding or holoenzyme assembly [3, 5].

Even less well understood are the structural properties of pseudoenzymes that allow them to carry out these functions. Bioinformatic analyses imply that most pseudoenzymes have evolved from ancestral cognate enzymes due to loss of one or more residues required for catalysis or co-factor binding [6, 7]. However, it is difficult to assess bioinformatically whether pseudoenzymes have acquired additional mutations beyond these catalytic site mutations that prevent their ancestral enzymatic function. Indeed, this question has only been experimentally addressed in a handful of studies [1]. Converting the degenerate glycine residue of the STYX pseudophosphatase to a catalytic cysteine readily restored its hydrolytic activity [8, 9]. In contrast, mutations to restore the degenerate sites of pseudokinases have had differential effects depending on the pseudokinase. When residues required for binding ATP were restored in the RYK pseudokinase, it re-gained kinase activity [10]. Conversely, the equivalent mutation in the HER3 pseudokinase failed to restore its kinase activity [11, 12]. Similarly, when a critical catalytic cysteine was restored to the DivL histidine pseudokinase of the bacterium *Caulobacter crescentus*, it did not regain kinase activity [13].

While the question of whether pseudoenzymes can be converted back into active enzymes has been experimentally tested for pseudophosphatases and pseudokinases, this question has not yet been examined for pseudoproteases. In this study, we attempted to resurrect the protease activity of two pseudoproteases, CspA and CspC, which play critical roles in the life cycle of *Clostridioides difficile*, a spore-forming bacterial pathogen that is the leading cause of nosocomial gastroenteritis worldwide [14, 15]. In the U.S. alone, *C. difficile* caused ∼225,000 infections and ∼13,000 deaths in 2017, leading to medical costs in excess of $1 billion [16]. Indeed, *C. difficile* has been classified as an urgent threat by the Centers for Disease Control because of its intrinsic resistance to antibiotics and the risk this poses to patients undergoing antimicrobial treatment [17].

*C. difficile* infections are transmitted by its incredibly resistant and infectious spore form. Because *C. difficile* is an obligate anaerobe, its vegetative cells cannot survive in the presence of oxygen [18, 19]. Thus, *C. difficile* infections begin when its metabolically dormant spores germinate in the gut of vertebrate hosts in response to bile acids [20]. Notably, unlike almost all other spore-formers studied to date, *C. difficile* senses bile acid germinants instead of nutrient germinants [21, 22]. The bile acid germinant signal is transduced by clostridial serine proteases known as the Csps [23-26], which are members of the subtilisin-like serine protease family members [27, 28] conserved in many clostridial species [29]. In these organisms, the three Csp proteins, CspA, CspB and CspC, participate in a signaling cascade that leads to the proteolytic activation of the SleC cortex hydrolase. Activated SleC then removes the protective cortex layer, which is essential for spores to exit dormancy [23, 30].

While CspA, CspB, and CspC are all active in *Clostridium perfringens* [28], *C. difficile* CspC and CspA carry substitutions in their catalytic triad that render them pseudoproteases [23, 24, 31]. CspC is thought to directly sense bile acid germinants [24] and integrate signals from cation and amino acid cogerminants that potentiate germination in the presence of bile acid germinants [32]. CspA has been proposed to function as the cation and amino acid co-germinant receptor [26] while also being necessary for CspC to be packaged into mature spores [25]. Thus, both CspC and CspA are required to activate the CspB protease, which subsequently cleaves the SleC cortex hydrolase [23]. CspB is a canonical subtilisin-like serine protease family member: it carries an intact catalytic triad consisting of aspartate, histidine, and serine residues and a long N-terminal prodomain that serves as an intramolecular chaperone to induce proper folding of the protease domain [27]. Like other active subtilases [27], once the protease domain of CspB adopts an active conformation, it undergoes autoprocessing to separate the prodomain from the protease domain. In contrast, CspA and CspC do not undergo autoprocessing in *C. difficile* [23, 25], presumably because they both carry mutations in their catalytic triad.

Interestingly, *cspA* and *cspB* are encoded in a single ORF, *cspBA*, such that the resulting fusion protein physically links the active CspB protease to the inactive CspA pseudoprotease. CspBA’s protease-pseudoprotease arrangement is largely conserved in the Peptostreptococcaceae family to which *C. difficile* belongs [25], with the CspB domain carrying an intact catalytic triad in all sequences examined and the CspA domain typically carrying at least one mutation in its catalytic triad ([25], **Figure 1A**). While the catalytic site mutations present in the CspA pseudoprotease vary in the Peptostreptococcaceae family, the degenerate site residues of CspC are strictly conserved within this family ([25], **Figure 1B**). In contrast, members of the Lachnospiraceae and Clostridiaceae families encode all three Csp proteins as individual proteases with intact catalytic triads. Accordingly, all three Csp proteins in *C. perfringens* undergo autoprocessing in mature spores [28], indicating that they are active proteases that can presumably process pro-SleC to active SleC cortex hydrolase in spores [33]. Based on these observations, it seems likely that the CspA and CspC pseudoproteases of the Peptostreptococcaceae family are the “odd ones out” among clostridial serine proteases (Csps) in their lack of catalytic activity.

**Figure 1.**
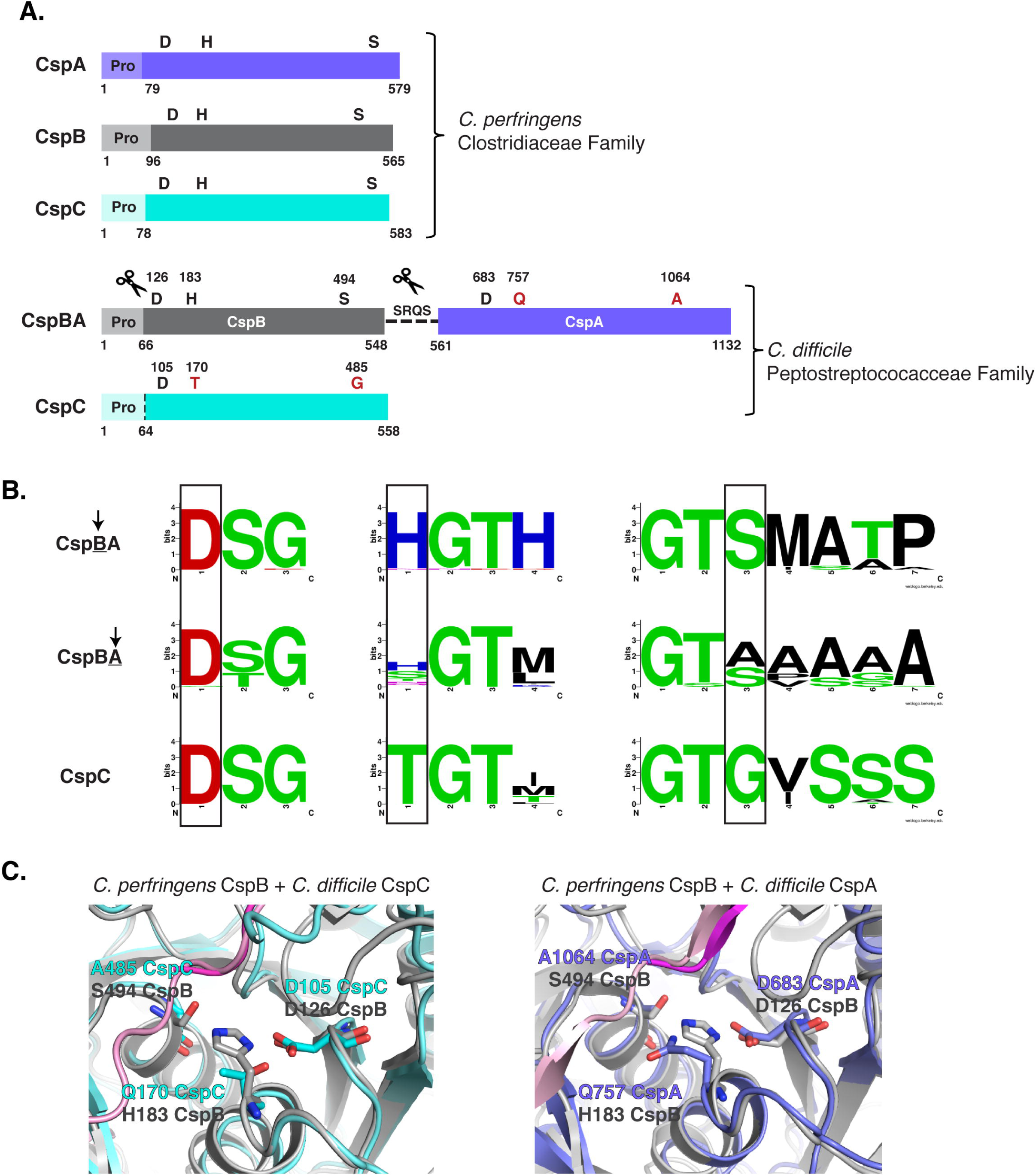
Csp family subtilisin-like serine proteases in the Clostridia. (A) Schematic of the active Csp proteases encoded by *C. perfringens*, CspA, CspB and CspC, compared to *Clostridiodes difficile* Csp proteins, where the active CspB protease is fused to an inactive CspA pseudoprotease domain, and CspC is also a pseudoprotease. “Pro” denotes the prodomain that functions as an intramolecular chaperone. Catalytic triad residues aspartic acid (D), histidine (H) and serine (S) are shown in black; degenerate site residues are shown in red (glutamine (Q), aspartic acid (A), threonine (T), and glycine (G). The scissor icon marks autoprocessing of Csp prodomains, whereas the dotted line indicates where the prodomain would be autoprocessed if the CspC pseudoprotease were to become active. The proposed YabG cleavage site around residues SQRS [26] is shown between CspB and CspA. (B) Sequence logos of the catalytic triad residue regions for CspBA and CspC of the Peptostreptococcaceae family. Regions shown correspond to the MEROPS protease database [43] definitions for the peptidase family S8A. Information regarding gene location and accession number for the proteins is included in the sequence logo analysis provided in Table S3. (C) Cartoon model of the active/degenerate site regions of either *C. perfringens* CspB (grey, PDB 4I0W) aligned to *C. difficile* CspC (cyan, PBD 6MW4) or CspB (grey, PDB 4I0W) aligned to *C. difficile* CspA iTasser model (periwinkle), active site region. The cleaved prodomain of CspB is shown in dark magenta compared to the uncleaved prodomain of CspC (pink) and CspA prodomain (light pink).

Interestingly, we recently showed that CspC’s degenerate site residues closely align with the catalytic triad of the active CspB protease in the X-ray crystal structure of CspC ([32], **Figure 1B**), leading us to query whether restoring an intact catalytic triad would be sufficient to convert CspC into an active protease capable of undergoing autoprocessing like other subtilisin-like serine proteases [27]. Since structural modeling of CspA also predicted close alignment of its degenerate site residues with CspB’s catalytic triad residues (**Figure 1B**), we tested the effect of restoring the catalytic triads of *C. difficile* CspC and CspA to gain insight into the evolution of these “zombie” proteins from active proteases and the structural requirements for their function in *C. difficile*.

## Experimental

### Bacterial strains and growth conditions

*C. difficile* strain construction was performed using 630Δ*erm*Δ*cspC*Δ*pyrE* [31] and 630Δ*erm*Δ*pyrE*-Δ*cspBA* as the parental strains via *pyrE*-based allele-coupled exchange (ACE [34]). This system allows for single-copy complementation of the Δ*cspC* and Δ*cspBA* parental mutants, respectively, from an ectopic locus. *C. difficile* strains are listed in **Table S1**. They were grown on brain heart infusion media (BHIS) supplemented with taurocholate (TA, 0.1% w/v; 1.9 mM), thiamphenicol (10-15 µg/mL), kanamycin (50 µg/mL), cefoxitin (8 µg/mL), and L-cysteine (0.1% w/v; 8.25 mM) as needed. Cultures were grown under anaerobic conditions at 37°C using a gas mixture containing 85% N_2_, 5% CO_2_, and 10% H_2_.

*Escherichia coli* strains for BL21(DE3)-based protein production and for HB101/pRK24-based conjugations are listed in **Table S1**. *E. coli* strains were grown shaking at 225 rpm in Luria-Bertani broth (LB) at 37 °C. The media was supplemented with ampicillin (50 µg/mL), chloramphenicol (20 µg/mL) or kanamycin (30 µg/mL) as indicated.

### *E. coli* strain construction

*E. coli* strains are listed in **Table S1** in the supplementary material. As previously described [32], the *cspC* complementation constructs were created using flanking primers, #2189 and 2242 (**Table S3**), in combination with internal primers encoding a given point mutation, Δ*cspBA* genomic DNA was used as the template. This resulted in *cspC* complementation constructs carrying 282 bp of the *cspBA* upstream region in addition to the Δ*cspBA* sequence and the intergenic region between *cspBA* and *cspC*. This extended construct was required to produce wild-type levels of CspC when expressing the constructs in the *pyrE* locus [31, 34]. For example, the T170H mutation was constructed using primer pair #2189 and 2355 to amplify a 5’ *cspC* complementation construct fragment encoding the T170H mutation at the 3’ end, while primer pair #2354 and 2242 were used to amplify a 3’ *cspC* complementation construct encoding the T170H mutation at the 5’ end. The individual 5’ and 3’ products were cloned into pMTL-YN1C digested with NotI/XhoI by Gibson assembly. In some cases, the two PCR products were used in a PCR SOE [35] prior to using Gibson assembly to clone the *cspC* construct into pMTL-YN1C digested with NotI and XhoI. The resulting plasmids were transformed into *E. coli* DH5α, confirmed by sequencing, and transformed into HB101/pRK24.

Similarly, for *cspBA* complementation constructs, each construct was designed with 126 bp of the Δ*cspC* sequence downstream of *cspBA* in order to fully complement the Δ*cspBA* mutant as previously described [31]. All primers used for strain construction are listed in **Table S3**. For example, the Q757H point mutation was introduced into the complementation constructs by using primer pair #2189 and #3041 to amplify the 5’ end, and #3040 and #2242 to amplify the 3’ end. The A1064S mutant was designed in the same way, but with primer pair #3042 and #3043 to introduce the point mutation. The 5’ and 3’ products containing the various mutations were cloned into pMTL-YN1C digested with NotI/XhoI and combined through Gibson assembly. Depending on the construct, some PCR products were combined by PCR SOE prior to using Gibson assembly. The resulting plasmids were transformed into *E. coli* DH5α, confirmed by sequencing, and transformed into HB101/pRK24.

To clone the construct encoding the *cspBA* prodomain trans-complementation construct, primer pair #2189 and 951 was used to amplify the 5’ fragment, and primer pair #950 and 2242 were used to amplify the 3’ fragment. In both cases, Δ*cspC* genomic DNA was used as a template as described previously [31]. The resulting two fragments were joined together using PCR SOE with primer pair #2189 and 2242, and the PCR SOE product was cloned into pMTL-YN1C digested with NotI and XhoI using Gibson assembly. A similar strategy was used to generate the *cspC* prodomain trans-complementation construct. Primers #2189 and 2553 were used to amplify the 5’ fragment, and primer pair #2552 and 2242 were used to amplify the 3’ fragment using Δ*cspBA* genomic DNA as a template. The fragments were joined together using SOE PCR with the primer pair #2189 and 2242, and the resulting SOE PCR product was cloned into pMTL-YN1C digested with NotI and XhoI using Gibson assembly [32].

To generate the recombinant protein expression constructs for producing CspC-His_6_ variants, primer pair #1128 and 1129 was used to amplify a codon-optimized version of *cspC* using pJS148 as the template (a kind gift from Joseph Sorg) as previously described [32]. The resulting PCR product was digested with NdeI and XhoI and ligated into pET22b cut with the same enzymes. The G457R variant was cloned using a similar procedure except that primer pair #1128 and 1361 and primer pair #1360 and 1129 were used to PCR the 5’ and 3’ fragments encoding the G457R mutation. The resulting PCR products were joined together using PCR SOE and flanking primer pair #1128 and 1129.

The remaining constructs encoding *cspC* codon-optimized variants for expression using pET22b were cloned using Gibson assembly. Flanking primer pair #2311 and 2312 were used to generate PCR products when used in combination with the internal primers encoding the point mutations. The resulting PCR products were cloned into pET22b digested with NdeI and XhoI using Gibson assembly. PCR SOE was sometimes used to join the two 5’ and 3’ fragments prior to Gibson assembly into pET22b. Similarly, the CspBA-His_6_ recombinant protein expression constructs, were constructed using primers #3034 and 3035 to create a codon-optimized version of *cspBA*. The Q757H and A1064S point mutations were introduced using primer pairs, #3036 and 3037, and #3038 and 3039, respectively. The resulting PCR products were digested with NcoI/XhoI, ligated into pET28a, and transformed into BL21.

To generate the recombinant protein expression constructs for producing CspBA-His_6_ variants, primer pair #1505 and 1529 was used to amplify a codon-optimized version of *cspB* using a plasmid template (a kind gift from Joseph Sorg). The resulting PCR product was digested with NcoI and HindIII and ligated into pET28a digested with the same enzymes. Codon-optimized *cspA* was then amplified using primer pair #1507 and 1508 using another plasmid template from Joseph Sorg. The resulting PCR product was used as the template for a second PCR using primer pair #1530 and 1508. This PCR product was digested with HindIII and XhoI and then ligated into pET28a-*cspB* CO digested with the same enzymes.

### Protein purification for Recombinant *E. coli* Analyses

*E. coli* BL21(DE3) strains listen in **Table S2** were used to produce and purify codon-optimized *cspC* variants and *cspBA* variants as previously described [23]. Briefly, cultures were grown to mid-log phase in 2YT (5 g NaCl, 10 g yeast extract, and 15 g tryptone per liter). When cultures reached an OD_600_ ∼0.8, 250 µM isopropyl-β-D-1-thiogalactopyranoside (IPTG) was added to induce expression of *cspC*. Cultures were then grown overnight at 18°C. The cells were pelleted, resuspended in lysis buffer (500 mM NaCl, 50 mM Tris [pH 7.5], 15 mM imidazole, 10% [vol/vol] glycerol), flash frozen in liquid nitrogen, thawed and finally sonicated. The insoluble material was pelleted, and the soluble fraction was incubated with Ni-NTA agarose beads (5 Prime) for 3 hrs, and eluted using high-imidazole buffer (500 mM NaCl, 50 mM Tris [pH 7.5], 200 mM imidazole, 10% [vol/vol] glycerol) after nutating the sample for 5-10 min.

### *C. difficile* strain construction

Complementation strains were constructed using CDDM to select for recombination of the complementation construct into the *pyrE* locus by restoring uracil prototrophy [34], as previously described [36]. At least two independent clones from each complementation strain were phenotypically characterized.

### Sporulation

*C. difficile* strains were grown overnight on BHIS plates containing taurocholate (TA, 0.1% w/v, 1.9 mM). Liquid BHIS cultures were inoculated from the resulting colonies, which were grown to early stationary phase before being back-diluted 1:50 into BHIS. When the cultures reached an OD_600_ between 0.35 and 0.75, 120 µL of this culture were plated onto 70:30 agar plates and grown for 18-24 hours as previously described [37]. Sporulating cells were harvested into phosphate-buffered saline (PBS), and cells were visualized by phase-contrast microscopy [38].

### Spore purification

Sporulation was induced on 70:30 agar plates for 2-3 days as described above, and spores were purified as previously described [30]. Briefly, the samples were harvested into sterile water at 4°C. The samples were washed 6-7 times in 1 mL of ice-cold water per every 2 plates and incubated overnight in water at 4°C. The samples were then pelleted and incubated with DNase I (New England Biolabs) at 37 °C for 60 minutes. Finally, samples were purified on a 20%:50% HistoDenz (Sigma Aldrich) gradient and washed 2-3 more times in water. Spore purity was assessed using phase-contrast microscopy (>95% pure). The optical density of the spore stock was measured at OD_600_, and spores were stored in water at 4 °C.

### Germination assay

As previously described [30], germination for each strain used the equivalent of 0.35 OD_600_ units, which corresponds to ∼1 × 10^7^ spores. The proper number of spores was resuspended in 100 µl of water,10 µL of this mixture was serially diluted in PBS, and the resulting dilutions were plated on BHIS-TA. Colonies arising from germinated spores were enumerated at 18-24 hrs. Germination efficiencies were calculated using mean CFUs produced by spores for a given strain relative to the mean CFUs produced by wild type. Analyses were based on at least three technical replicates performed on two independent spore preparations (i.e. two biological replicates). Statistical significance was determined by performing a one-way analysis of variance (ANOVA) on natural log-transformed data using Tukey’s test.

### OD_600_ kinetics assay

As previously described [39], approximately 1.5 × 10^7^ spores (0.48 OD_600_ unit) were resuspended in BHIS to a total volume of 1.1 mL. The sample was divided in two: 540 μl was added to a cuvette containing 60 μl of 10% taurocholate, while the other sample was added to a cuvette containing 60 μl of water, as a control. The samples were mixed, and the OD_600_ was measured every 3 min for 90 min. Statistical significance was determined by performing a twoway analysis of variance (ANOVA) using Tukey’s test.

### Western blot analysis

Samples for western blot analysis were prepared as previously described [40]. Briefly, sporulating cell pellets were resuspended in 100 µL of PBS, and 50 µL samples were removed and freeze-thawed for three cycles. The samples were resuspended in 100 µL EBB buffer (8 M urea, 2 M thiourea, 4% (w/v) SDS, 2% (v/v) β-mercaptoethanol), boiled for 20 min, pelleted, resuspended in the same volume. Subsequently, 7 uL of sample buffer was added to stain samples with bromophenol blue. *C. difficile* spores (∼1 × 10^7^) were resuspended in 50 µL EBB buffer and processed similarly. The samples were resolved by 7.5% (for sporulating cell analyses of CspBA and CspC) or 12% SDS-PAGE gels. After, the gels were transferred to Millipore Immobilon-FL PVDF membranes and were blocked in Odyssey Blocking Buffer [36] for 30 mins with 0.1% (v/v) Tween 20. Blots were incubated with rabbit polyclonal anti-CspB [23], anti-CspA (a generous gift from Joe Sorg, Texax A&M University, [26]) or anti-CotA antibodies and/or mouse polyclonal anti-SleC [23], anti-CspC [25], or anti-SpoIVA antibodies [41]. Additionally, western blotting for recombinant protein samples were blotted with commercial antibody, mouse monoclonal anti-penta-His (ThermoScientific). The anti-CspB, anti-CspC, anti-SpoIVA antibodies were used at 1:2500 dilutions, the anti-SleC antibody was used at a 1:5000 dilution, and the anti-pentaHis, anti-CotA, and anti-CspA antibodies were used at a 1:1000 dilution. IRDye 680CW and 800CW infrared dye-conjugated secondary antibodies were used at 1:20,000 dilutions. The Odyssey LiCor CLx was used to detect secondary antibody infrared fluorescence emissions. All blots shown are representative of analyses performed on two independent spore preparations.

### Protein Modeling

Multiple protein model predictions were used to analyze the CspA structure. Specifically, we analyzed several predictions by I-TASSER (Iterative Threading ASSEmbly Refinement) [42]. All prediction models were downloaded as PDB files and were viewed using PyMol. The CspA sequence used was taken from the *cspBA* gene, starting at codon 560 which has been predicted to encode the YabG cleavage site [26]. CspA predictions were aligned with the RCSB PDB files of the CspB (PDB 4I0W) protease and CspC pseudoprotease (PDB 6MW4).

### Protein Sequence Analysis

Protein sequences were obtained from NCBI protein by searching for homologous sequences to CspBA and CspC, respectively, in *Clostridioides difficile* strain 630 filtering only for species within the Peptostreptococcaceae family. The algorithm “PSI-BLAST” was used to identify distant relatives of the proteins of interest. Homologs that had >95% query cover were selected for analysis, and for CspC homologs only, sequences which additionally had >55% identity were selected to avoid redundancy with CspA or CspB individual homologs. For all analyses only the first 3 organisms from each species were selected, in order to avoid skewing of the data based on the most sequenced organisms. The selected sequences were analyzed using MacVector and aligned with the ClustalW algorithm. The regions surrounding the catalytic residues were selected based on previous analyses [25] using the MEROPS protease database [43]active site definitions for peptidase family S8A. Information regarding accession numbers for selected homologs is provided in **Table S3** and **Table S4**.

## Results

### Restoring CspC’s catalytic triad disrupts protein folding and abrogates its function

To determine if protease activity could be resurrected in the CspC pseudoprotease, we restored the degenerate residues of CspC’s catalytic triad (**Figure 1A**) individually and in combination. To this end, we generated strains producing CspC variants carrying the following amino acid substitutions: threonine 170 to histidine (CspC_T170H_), glycine 485 to serine (CspC_G485S_), and T170H-G485S (referred to as CspC_2xcat_). The constructs encoding these substitutions were integrated into the *pyrE* locus of a previously characterized in-frame *cspC* deletion mutant [31] using allele-coupled exchange [34]. These constructs, along with all other *cspC* constructs analyzed in this manuscript, were expressed from the native *cspBA-cspC* promoter as described previously [31].

To assess whether the T170H-G485S substitutions in CspC activated the autoprocessing activity characteristic of subtilisin-like serine proteases [27, 44], we analyzed CspC processing in sporulating cells using western blotting. Rather than restoring autoprocessing activity, mutations in the degenerate sites decreased CspC levels in sporulating cell lysates: no CspC was detectable in lysates of the double mutant (*2xcat*) strain (**Figure 2B**), and CspC levels were markedly diminished in the *G485S* mutant and slightly reduced in the *T170H* mutant. Taken together, mutations in CspC’s degenerate residues reduce CspC production and/or stability in sporulating cells. In contrast, CspBA levels were unaffected in the mutant strains (**Figure 2B**), consistent with the observation that CspC does not affect CspBA levels [31].

**Figure 2:**
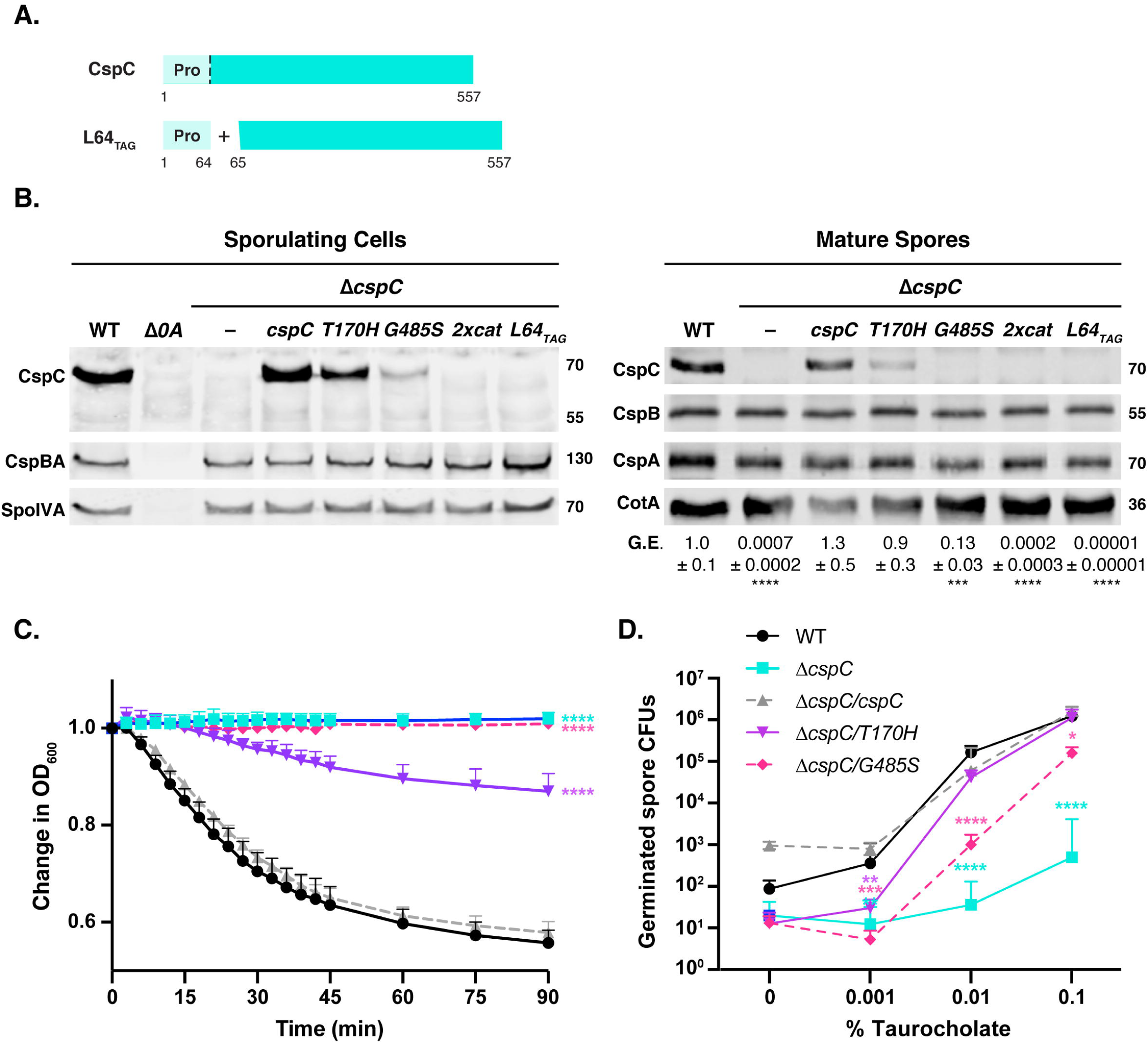
Restoring active site residues to CspC appears to result in its destabilization. (A) Schematic of wild-type CspC and a construct encoding the prodomain of CspC *in trans* (L64-TAG). The dotted line indicates where the prodomain would be autoprocessed if the CspC pseudoprotease were to become active. “Pro” denotes the prodomain. L64-TAG encodes a variant in which the CspC prodomain is produced *in trans* from the remainder of CspC through the introduction of a stop codon after codon 64 and a ribosome binding site and start codon before codon 65. (B) Western blot analyses of CspB(A) and CspC in sporulating cells and purified spores from wild type 630Δ*erm*-p, Δ*cspC*, and Δ*cspC* complemented with either wild-type *cspC* or the L64-TAG *cspC* trans-complementation variant. *2xcat* refers to a complementation construct containing both the T170H and G485S point mutations. Δ*spo0A* (Δ*0A*) was used as a negative control for sporulating cells. SpoIVA was used as a loading control for sporulating cells, while CotA was used as a loading control for purified spores. An anti-CspB antibody was used to detect full-length CspBA in sporulating cells, whereas CspB is detected in purified spores [23]. The germination efficiency of spores from the indicated strains plated on BHIS media containing 0.1% taurocholate is also shown relative to wild type. The mean and standard deviations shown are based on multiple technical replicates performed on two independent spore purifications for purified spores. Statistical significance relative to wild type was determined using a one-way ANOVA and Tukey’s test. (C) Change in the OD_600_ in response to germinant of CspC catalytic mutant spores relative to wild-type spores. Δ*cspC* mutant spores serve as a negative control. *2xcat* mutant spores are not shown as they behaved similarly to the Δ*cspC* spores in the less sensitive plate-based assay. Purified spores were resuspended in BHIS, and germination was induced by adding taurocholate (1% final concentration). The ratio of the OD_600_ of each strain at a given time point relative to the OD_600_ at time zero is plotted. The mean of three assays from at least 2 independent spore preparations are shown. The error bars indicate the standard deviation for each time point measured. Statistical significance relative to wild type was determined using a two-way ANOVA and Tukey’s test. (D) Germinant sensitivity of CspC catalytic mutant spores compared to wild type. Spores were plated on BHIS containing increasing concentrations of taurocholate. The number of colony forming units (CFUs) produced by germinating spores is shown. The mean and standard deviations shown are based on multiple replicates performed on two independent spore purifications. Statistical significance relative to wild type was determined using a one-way ANOVA and Tukey’s test. **** p < 0.0001, *** p < 0.001, ** p < 0.01.

To determine whether the reduced CspC protein levels in the degenerate site mutants were due to general protein folding defects, we cloned the mutant alleles into recombinant protein expression vectors and assessed His-tagged CspC variant levels and autoprocesssing in *E. coli.* We also assessed their purification efficiency from the soluble fraction of *E. coli* lysates. As a control, we cloned the *cspC*_*G171R*_ allele, which abrogates germination in *C. difficile*, [24] likely by disrupting CspC folding due to steric hindrance (**Figure 3A**, [32]). Recombinant CspC_2xcat_ did not undergo autoprocessing when produced in *E. coli* (**Figure 3B**, Induced fraction), indicating that restoring the catalytic triad does not reconstitute CspC protease activity. However, when recombinant CspC_2xcat_ was purified from the soluble fraction, markedly less CspC_2xcat_ was obtained relative to wild-type CspC-His_6_ (Elution fraction, **Figure 3B**) even though CspC_2xcat_ was observed at wild-type levels following IPTG induction (Induced fraction, **Figure 3B**). The purification yields for the single degenerate site variants were similarly reduced relative to wild-type CspC despite their wild-type induction levels in *E. coli*, and the predicted protein folding mutant, *G171R*, yielded the lowest amount of soluble purified CspC. Unfortunately, it was not possible to assess whether the mutant alleles reduced the solubility of CspC, because wild-type CspC was largely detected in the insoluble fraction in *E. coli* (data not shown).

**Figure 3.**
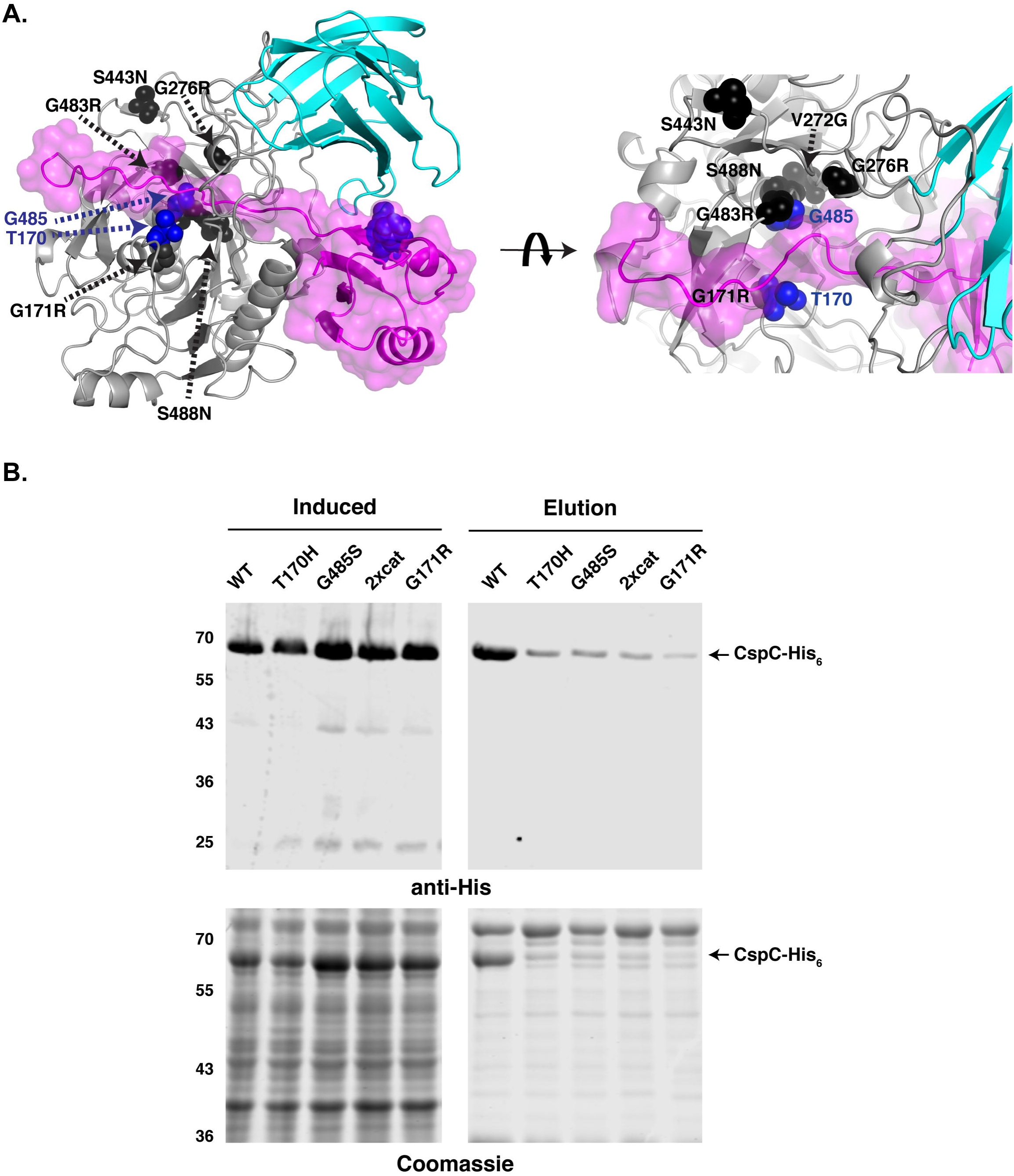
Restoring CspC’s catalytic triad appears to impair protein folding in *E. coli*. (A) CspC space fill model with jelly roll domain in cyan, prodomain in pink and subtilase domain in grey. Residues identified as being required for *C. difficile* spore germination by Francis et al. [24] in a genetic screen are shown in black. The S443N mutation was identified in combination with V272G. The degenerate site residues Thr170 and Gly485 are shown in blue. (B) Purification of CspC-His_6_ variants from the soluble fraction. G171R was included because this mutation had been predicted to destabilize CspC by steric occlusion [21, 32]. Cultures expressing the *cspC* variants were induced with IPTG overnight at 18°C, and aliquots were removed for analysis of the “induced” fraction. Cultures were harvested, and cells were lysed using sonication. Following a high-speed centrifugation, the cleared lysate containing soluble proteins was incubated with Ni^2+^-NTA agarose beads. CspC-His_6_ variants were eluted from the beads using imidazole (elution fraction). Equivalent volumes of samples were resolved by SDS-PAGE and analyzed by western blotting (top) and Coomassie staining (bottom).

### Degenerate site mutations decrease germination rates and germinant sensitivity

To evaluate how the CspC degenerate site mutations impacted CspC function during spore germination, we measured the ability of these mutant alleles to complement Δ*cspC*’s germination defect. Purified spores from wild-type, Δ*cspC*, Δ*cspC*/*cspC*, and the degenerate site mutant complementation strains were plated on rich media containing 0.1% taurocholate germinant, and the number of colony forming units (CFUs) that arose from germinating spores relative to wild type was determined. The *cspC*_*T170H*_ allele did not affect germination relative to wild type even though CspC_T170H_ protein levels were visibly decreased in western blot analyses of purified spores (**Figure 2B**). The *cspC*_*G485S*_ allele resulted in only a ∼10-fold defect in germination efficiency despite producing almost undetectable levels of CspC_G485S_ protein. In contrast, the *2xcat* double mutant exhibited a germination defect (∼1,000-fold) equivalent to that of the parental Δ*cspC* strain, consistent with the absence of detectable CspC_2xcat_ in sporulating cells and purified spores (**Figure 2B**). Low levels of “spontaneous” germination were observed in the Δ*cspC* spores, similar to previous analyses of other germinant receptor mutants [31, 36, 45, 46]. These results indicate that relatively little CspC is needed to allow *C. difficile* spores to germinate.

Although the *G485S* mutant exhibited only an ∼10-fold germination defect when spores were plated on rich media containing 0.1% germinant, G485S colonies arose more slowly than colonies derived from wild-type spores. To test whether the G485S and T170H degenerate site mutations affected the rate of germination, we used an optical density-based germination assay. This assay measures the decrease in optical density of a population of germinating spores over time due to cortex hydrolysis and core hydration [39]. While the optical density of wild-type spores decreased by ∼40%, the optical density of the G485S mutant did not appreciably change over the assay period, similar to the Δ*cspC* strain (**Figure 2C**, p < 0.0001). Surprisingly, the T170H mutant germinated more slowly relative to wild type (p < 0.0001) even though *cspC*_T170H_ did not exhibit a germination defect in the plate-based CFU assay. Taken together, these results indicate that single mutations in CspC’s degenerate active site impair folding and/or decrease CspC stability in sporulating cells. These decreased CspC levels correlate with slower spore germination rates. Furthermore, rather than conferring autoprocessing activity to CspC, restoring its full catalytic triad appears to cause protein misfolding and reduces CspC levels in spores.

We next wondered whether the reduced CspC levels in the *T170H* and *G485S* mutant spores would decrease their germinant sensitivity. We had previously determined that spores lacking GerG have decreased CspB, CspA, and CspC levels, which correlated with diminished responsiveness to germinant [36]. Importantly, unlike *gerG* mutant spores, the degenerate site mutants carry wild-type levels of CspB and CspA in purified spores (**Figure 2B**). Thus, germination defects in CspC degenerate site mutants can be attributed to impaired CspC function and/or decreased protein levels rather than changes in CspB or CspA levels. To measure germinant sensitivity, we plated *cspC* mutant spores on rich media containing varying concentrations of taurocholate germinant. On plates containing 0.001% taurocholate, *T170H* and *G485S* mutant spores germinated to a similar extent as Δ*cspC* spores (p ≤ 0.005, **Figure 2D**). However, on plates with 0.01% taurocholate, *T170H* mutant spores germinated to near wild-type levels, whereas *G485S* spores exhibited an ∼100-fold decrease relative to wild type (p < 0.0001). At the highest concentration of germinant tested (0.1% taurocholate), *T170H* mutant spores were indistinguishable from wild-type spores, and the *G485S* mutant spores exhibited an ∼10-fold decrease in CFUs (**Figure 2D**, p < 0.0001) consistent with our prior findings (**Figure 2B**). Thus, decreased CspC protein levels and/or function in the degenerate site mutants impairs germinant sensing even when CspB and CspA are present at wild-type levels, consistent with CspC’s proposed role as the germinant receptor [24].

### The CspC prodomain cannot function in *trans*

Since our attempts to “resurrect” CspC’s active site appeared to destabilize the protein, we wondered whether we could bypass the autoprocessing event by producing the CspC prodomain separate from the CspC subtilase domain. As mentioned earlier, subtilisin-like serine proteases use their long N-terminal prodomain as an intramolecular chaperone to promote folding of the subtilase domain into an active conformation [44]. The prodomains of other subtilisin-like serine proteases (including CspB in *C. difficile*) can perform this chaperone function *in trans* [23, 27, 44]. We thus tested whether *C. difficile* CspC’s prodomain could function as a chaperone *in trans*, even though CspC normally does not undergo autoprocessing. To this end, we generated a complementation construct (L64_TAG_) that produced the prodomain (residues 1-64) separate from the remainder of the CspC protein (residues 65-557, **Figure 2A**). CspC was undetectable in the *L64*_*TAG*_ mutant in western blot analyses of sporulating cells or purified spores (**Figure 2B**), suggesting that the CspC prodomain cannot function *in trans* unlike CspB [23] and other subtilisin-like serine proteases [27].

However, an important caveat to our prior finding that the CspB prodomain could function *in trans* was that these studies used plasmid over-expression [23]. Given that we recently determined that plasmid-based *cspBA-cspC* over-expression constructs can cause experimental artifacts [25, 31], we tested whether chromosomally encoding the CspB prodomain *in trans* would allow for complementation of a *cspBA* deletion strain (*Q66*_*TAG*_, **Figure S1A**). CspBA was detectable in sporulating cells of the *Q66*_*TAG*_ complementation strain, albeit at reduced levels relative to wild type and the wild-type *cspBA* complementation strain presumably because the chaperone activity of an intramolecular chaperone is more efficient than an intermolecular chaperone (**Figure S1B**). Regardless, these results indicate that the CspB protease can still fold properly when its prodomain is supplied *in trans* even when *cspBA*_*Q66-TAG*_ is expressed from the chromosome rather than a plasmid.

To assess how the decreased CspBA levels in the Q66_TAG_ complementation strain would affect CspB, CspA, and CspC levels in mature Q66_TAG_ spores (**Figure S1B**), we analyzed the levels of these proteins in purified spores by western blotting. Consistent with our prior report that CspB and CspA are needed to incorporate and/or stabilize CspC in mature spores [25, 31], reduced levels of CspB, CspA, and CspC were observed in Q66_TAG_ spores. Importantly, the Q66_TAG_ construct largely complemented the germination defect of the parental Δ*cspBA* strain, increasing the number of germinating spores by 1000-fold relative to Δ*cspBA* (p < 0.0001) and only 2-fold lower than wild-type spores (p < 0.01). We next tested whether the reduced levels of Csp proteins affected germinant sensitivity. Surprisingly, the greatly reduced levels of all three Csp proteins in Q66_TAG_ spores resulted in only a ∼2-fold defect in spore germination relative to wild type on 0.1% taurocholate plates and only a ∼4-fold decrease at the lowest concentration of taurocholate tested (0.001%, **Figure S1C**, p < 0.005). This result suggests that Csp proteins are present in excess of what is needed to respond to germinant signals.

### Resurrection of CspA’s active site does not restore enzymatic function

Our finding that the CspC pseudoprotease could not be resurrected by restoring its catalytic triad was perhaps not entirely surprising given that the degenerate site residues, Thr170 and Gly485, are strictly conserved across the Peptostreptococcaceae family ([25], **Figure 1B**). In contrast, the degenerate site mutations in CspA’s active site region are not strictly conserved, with Peptostreptococcaceae family members encoding CspA domains that carry either one or two mutations in residues of the catalytic triad and occasionally none at all (**Figure 1**) within the context of CspBA fusion proteins. Since some Peptostreptococcaceae variants appear to encode active CspA domains, *C. difficile* CspA might tolerate degenerate site mutations more readily than CspC. Despite the sequence variation observed around the CspA degenerate site among Peptostreptococcaceae family members, modeling this degenerate site using iTasser [42] indicated that the CspA pseudoprotease should have high structural homology to the CspC pseudoprotease (**Figure 1C**) and CspB proteases. Thus, we tested whether CspA could be converted back into an active protease by using allele-coupled exchange [34] to restore the catalytic triad of CspA. In particular, we cloned complementation constructs encoding amino acid substitutions of glutamine 757 to histidine (Q757H) and alanine 1064 to serine (A1064S) both individually and in combination. In contrast with the CspC degenerate site mutants (**Figure 2**), CspA degenerate site mutants did not affect CspBA levels in sporulating cells even in the mutant carrying an intact catalytic triad (*BA*_*2xcat*_, **Figure 4A**). It should be noted that in sporulating cells, CspBA is the predominant form observed, while CspA is separated from CspB in mature spores through the action of the YabG protease [23]. Notably, the degenerate site mutations did not affect CspBA function, since all three degenerate site mutants made functional, heat-resistant spores at wild-type levels **(Figure 4A)**. Nevertheless, despite folding normally, the *cspBA 2xcat* mutant failed to undergo autoprocessing, indicating that restoring the catalytic triad was not sufficient to convert CspA into an active protease (**Figure 4A**).

**Figure 4.**
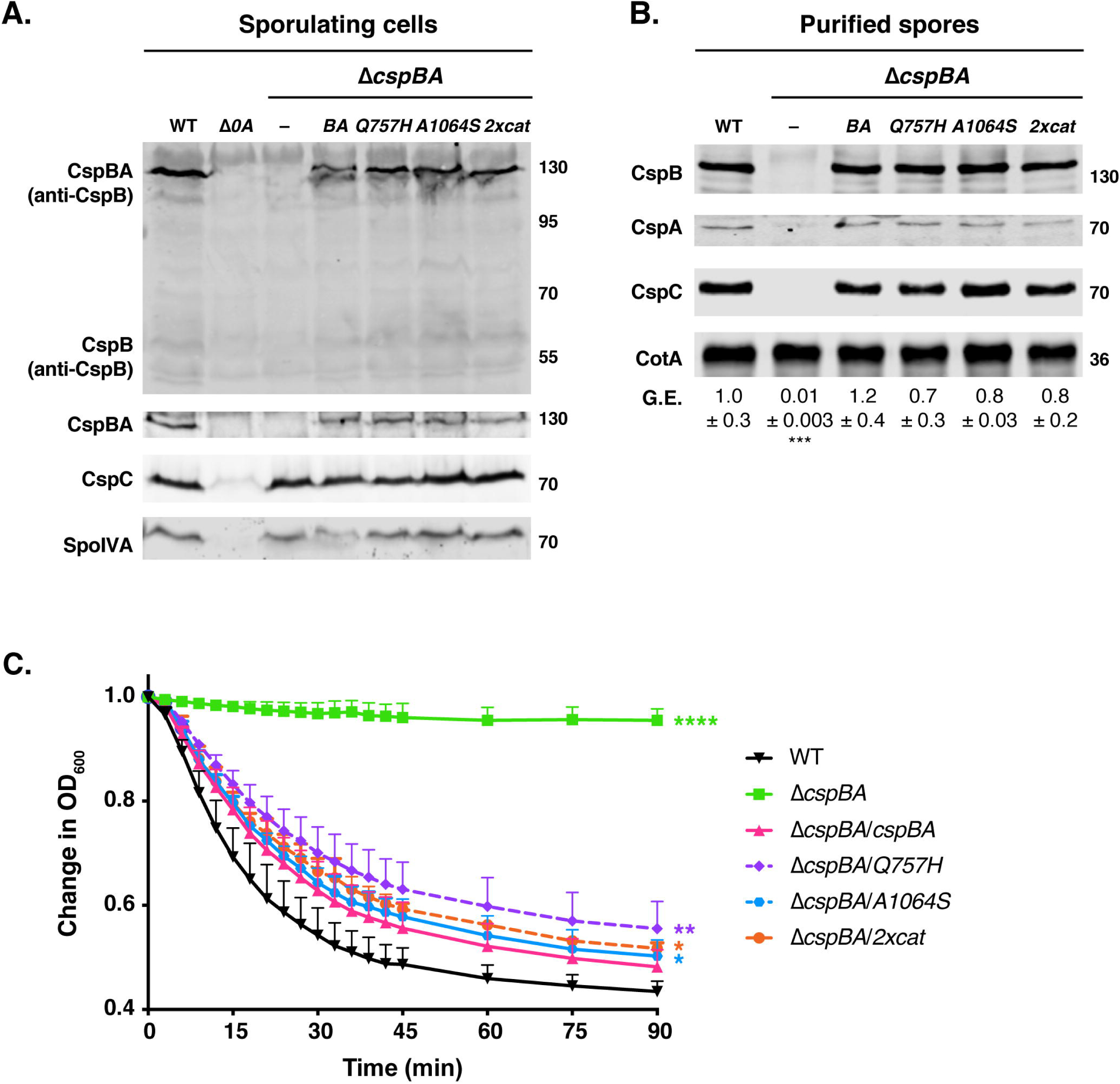
Restoring CspA’s catalytic triad in *C. difficile* does not resurrect its protease activity or impact CspBA levels or function. (A) Western blot analyses of CspB(A) and CspC in sporulating cells from wild type 630Δ*erm*-p, Δ*cspBA*, and Δ*cspBA* complemented with either wild-type *cspBA* or the *cspBA* catalytic variant (*2xcat*). CspBA is shown with anti-CspB antibody and anti-CspA antibody. Δ*spo0A* (Δ*0A*) was used as a negative control for sporulating cells. Anti-SpoIVA was used as a loading control for sporulating cells. (B) Western blot analyses of CspA, CspB and CspC in purified spores from wild type 630Δ*erm*-p, Δ*cspBA*, and Δ*cspBA* complemented with either wild-type *cspBA* or the *cspBA* catalytic variant (*2xcat*). CotA was used as a loading control for purified spores. The germination efficiency of spores from the indicated strains plated on BHIS media containing 0.1% taurocholate is also shown relative to wild type. The mean and standard deviations shown are based on multiple technical replicates performed on two independent spore purifications. Statistical significance relative to wild type was determined using a one-way ANOVA and Tukey’s test. (C) Change in the OD_600_ in response to germinant of CspBA catalytic mutant spores relative to wild-type spores. Δ*cspBA* mutant spores served as a negative control. Purified spores were resuspended in BHIS, and germination was induced by adding taurocholate (1% final concentration). The ratio of the OD_600_ of each strain at a given time point relative to the OD_600_ at time zero is plotted. The mean of three independent assays from at least 2 independent spore preparations are shown. The error bars indicate the standard deviation for each time point measured. Statistical significance relative to wild type was determined using a two-way ANOVA and Tukey’s test. **** p < 0.0001, *** p < 0.001, ** p < 0.01

Since CspB and CspA are separated from each other in mature spores [25], we considered the possibility that autoprocessing of CspA in the *BA*_*2xcat*_ double mutant might occur during spore maturation after YabG-mediated cleavage, so we analyzed the sizes and amount of CspB, CspA, and CspC in purified spores by western blotting. CspA_2xcat_ was detected at wild-type levels and at the expected size in degenerate site mutant spores (**Figure 4B**), confirming that CspA still does not undergo autoprocessing even if its catalytic triad is intact. CspB and CspC sizes and levels were similarly unaffected in the CspA degenerate site mutants (**Figure 4B**), and the mutant spores germinated with similar efficiency as wild type on rich media plates containing taurocholate germinant (**Figure 4B**). When measuring germination in the optical density-based germination assay, the mutant spores also exhibited similar drops in optical density relative to the wild-type *cspBA* complementation strain (Δ*cspBA*/*cspBA*), although they germinated slightly slower than wild-type spores in this assay (**Figure 4C**). Taken together, these results indicate that factors beyond CspA’s degenerate catalytic triad prevent the CspA pseudoprotease from functioning as an active protease and that CspA can tolerate mutations in its degenerate site region more readily than CspC.

While the inability to resurrect CspA’s protease activity likely reflects structural differences between the *C. difficile* CspA pseudoprotease and active Csp proteins in other clostridial organisms, an unknown inhibitory factor in *C. difficile* could prevent CspA_2xcat_ from acquiring autoprocessing activity. To test this possibility, we cloned codon-optimized degenerate site mutant alleles of *cspBA* into vectors for IPTG-inducible recombinant protein production in *E. coli*. We then measured CspBA-His_6_ variant production and purification levels in *E. coli*. Constructs encoding full-length CspBA were generated to reflect the form of CspA first produced in *C. difficile* sporulating cells, and producing CspA in the absence of CspB appears to destabilize CspA in sporulating *C. difficile* cells [31] and renders CspA largely insoluble in *E. coli* (data not shown).

CspBA_2xcat_-His_6_ was produced and purified at wild-type levels based on Coomassie staining (**Figure 5A**) and western blotting analyses (**Figure 5B**) in contrast with CspC_2xcat_-His_6_. However, similar to CspC_2xcat_-His_6,_ no CspA autoprocessing was observed in CspBA_2xcat_ (**Figure 5**), which would have led to CspA becoming separated from CspB. Unexpectedly, marked decreases in full-length CspBA_Q757H_ were observed in the IPTG-induced fraction compared to wild-type and the other CspBA variants. CspBA_Q757H_ appeared to be susceptible to protease cleavage in *E. coli* based on its altered banding pattern in the elution fraction relative to wild-type and the other CspBA variants (**Figure 5**). However, since the Q757H mutation did not affect CspBA size or function in *C. difficile* (**Figure 4a**), the point mutation likely alters the conformation of CspBA such that it is more susceptible to proteases present in *E. coli* but not *C. difficile*. Taken together, this data demonstrates that intrinsic structural features within the CspA pseudoprotease prevent it from being converted to a functional enzyme, even when its catalytic triad is restored.

**Figure 5.**
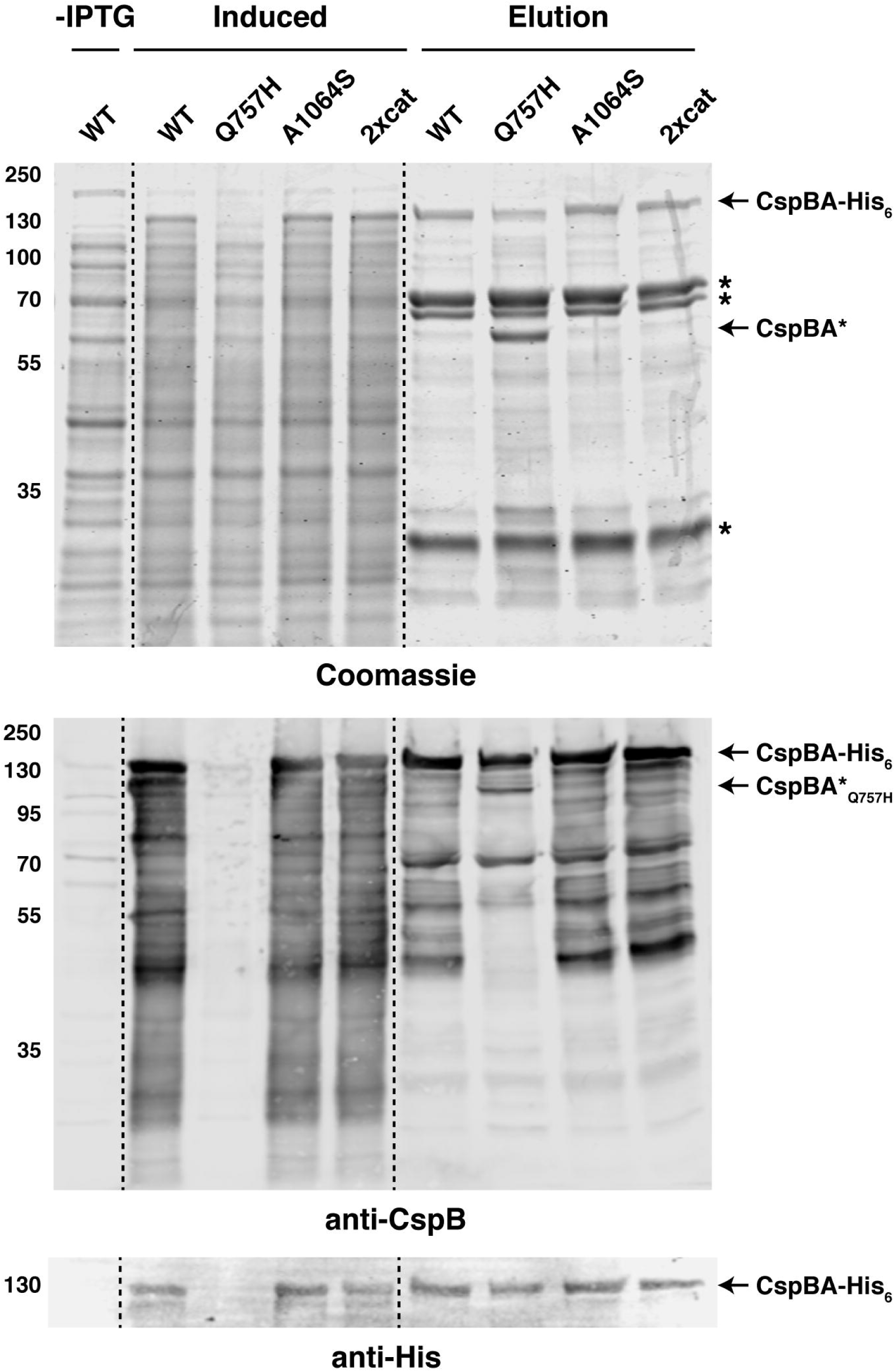
Resurrection of the CspA active site does not restore protease activity in *E. coli*. (A) Purification of CspBA variants from the soluble fraction. Cultures expressing the *cspBA* variants were induced with IPTG overnight at 18°C, and aliquots were removed for analysis of the “induced” fraction compared to the “uninduced” fraction prior to the addition of IPTG (-IPTG). Cultures were harvested, and cells were lysed using sonication. Following a high-speed centrifugation, the cleared lysate containing soluble proteins was incubated with Ni^2+^-NTA agarose beads. CspBA-His_6_ variants were eluted from the beads using imidazole (elution fraction). Samples were resolved by SDS-PAGE and analyzed by Coomassie staining (top) and western blotting (bottom). Non-specific proteins pulled-down with the Ni-NTA beads are marked with asterisks; a truncated CspBA Q757H variant is also marked. Anti-CspB antibody was used to detect full-length CspBA. Anti-His antibody was used to detect CspBA-His_6_ variants.

## Discussion

Pseudoenzymes have increasingly been recognized as crucial regulators of numerous cellular processes despite their lack of enzymatic activity [5]. These catalytically inactive variants most frequently arise from gene duplications followed by catalytic site inactivating mutations that then enable the pseudoenzyme to acquire a new non-enzymatic function [6, 7]. The extent to which a given pseudoenzyme’s function has diverged from its ancestral enzymatic function varies for the relatively small number of pseudoenzymes in which this question has been examined, with mutagenesis enabling the revival of enzymatic activity in some but not all pseudoenzymes [1]. Here, we assessed whether the CspA and CspC pseudoproteases of *C. difficile* could be converted to active proteases through the restoration of their catalytic triads. We also determined whether mutation of their degenerate site resides impacted CspC and/or CspA function during *C. difficile* spore germination.

Our mutational analyses revealed that restoring the active sites of the pseudoenzymes, CspC and CspA, did <u>not</u> resurrect protease activity in either pseudoenzyme regardless of whether the degenerate site mutants (*2xcat*) were produced in *C. difficile* or recombinantly in *E. coli*. Thus, the CspC and CspA pseudoproteases have both acquired features that prevent protease activity beyond their catalytic site mutations. Interestingly, the CspC and CspA pseudoproteases differentially tolerated changes to their degenerate site residues. Restoring CspC’s catalytic triad appeared to disrupt protein folding in *E. coli* (**Figure 3**) and *C. difficile* (**Figure 2**), whereas the equivalent mutations in CspA did not impact CspA folding or function in either organism (**Figures 4** & **5**). Given that these pseudoproteases tolerate changes in their degenerate sites to different degrees, our findings raise the question as to how CspC and CspA independently evolved to become pseudoproteases in *C. difficile* and other Peptostreptococcaceae family members.

In the case of CspC, loss of its catalytic site residues was likely critical to its evolution as a pseudoprotease. Although the crystal structure of CspC suggests that the substrate binding pocket can accommodate the catalytic triad residues **(Figure 1**, [32]), restoring these residues disrupts CspC folding (**Figure 3**). Analysis of the conservation of degenerate site residues in CspC homologs in the Peptostreptococcaceae family suggests that the chemical properties of its specific degenerate site residues, namely threonine 170 and glycine 485 in *C. difficile* CspC (**Figure 1**, [25]), are crucial for the structural integrity of Peptostreptococcaceae family CspC homologs.

Our prodomain transcomplementation experiments further revealed that additional changes beyond CspC’s degenerate site mutations appear to have been necessary for CspC to evolve its regulatory function during spore germination. Unlike CspB and many other subtilisin-like serine proteases whose subtilase domains still fold around their prodomains when supplied *in trans* (**Figure S1**, [23, 27, 44]), the prodomain of CspC lacked chaperone activity when supplied *in trans* (**Figure 2B**). The requirement for CspC’s prodomain to remain physically tethered to its subtilase domain may reflect the fact that the prodomain of *C. difficile* CspC is bound more tightly to its subtilase domain than the prodomain of *C. perfringens* CspB [23, 32]. For example, in *C. difficile* CspC, a “clamp” region holds the prodomain in place, while this feature is absent in other subtilisin-like proteases [32]. Lastly, loss-of-function *cspC* alleles identified in a prior genetic screen by Francis *et al.* [24] cluster to the degenerate site region and may prevent prodomain binding to the degenerate site region via steric occlusion (**Figure 3A**). Consistent with this interpretation, the *cspC*_*G171R*_ allele in recombinant CspC exhibited a substantially reduced purification efficiency from the soluble fraction than wild-type CspC-His_6_ and even relative to the other degenerate site mutants (**Figure 4**). This result strongly suggests that the loss-of-function CspC mutations identified in the germination mutant screen [24] disrupt CspC folding and lead to its destabilization in sporulating cells.

While CspC structure and function rely critically on the identity of its degenerate site residues, CspA structure and function were unaffected by mutations that restore the catalytic triad. This result suggests that loss of CspA’s catalytic residues was not an essential step in its evolution to become a pseudoprotease in *C. difficile*, in contrast with *C. difficile* CspC. While mutation of catalytic site residues is thought to be the most frequent driving force behind the evolution of new functions for pseudoenzymes, mutations that prevent substrate binding or catalysis have also been observed in some pseudoenzymes [6, 7]. It is likely that *C. difficile*’s CspA pseudoprotease domain has evolved analogous mutations that prevent it from cleaving its prodomain. It is possible that CspA could have evolved these differences relative to CspC because CspA plays an additional role in regulating CspC incorporation into spores [25, 31]. In the absence of a CspA crystal structure, the effect of these mutations remains unclear, but our results raise the important possibility that CspA domains in Peptostreptococcaeae family CspBA homologs with an intact catalytic triad (**Figure 1C**) may include mutations that occlude substrate binding and thus may actually be pseudoproteases.

A similar case of pseudoenzymes lacking catalytic activity despite retaining their active site residues has been observed in the iRhom family of proteins, which are widely conserved, inactive homologs of rhomboid proteases [6]. While most iRhom pseudoproteases lack either one or both catalytic dyad residues required for rhomboid protease activity, at least one iRhom family member carries an intact catalytic dyad yet lacks protease activity [47]. Notably, a distinguishing feature of iRhoms is the presence of a proline residue adjacent to the catalytic serine (or degenerate site residue) that is sufficient to prevent proteolytic activity in rhomboid proteases [47]. Whether a similar type of inactivating residue occurs in the CspA (or CspC) pseudoproteases in the Peptostreptococcaceae family remains to be determined. Regardless, it is worth noting that iRhom pseudoproteases carry additional structural features that distinguish iRhoms from rhomboid proteases beyond the catalytic site mutations [6], namely an extended cytoplasmic amino terminus and conserved cysteine-rich luminal loop domain.

Unlike iRhoms and rhomboid proteases, the CspC pseudoprotease exhibits a high degree of structural similarity to the CspB protease, with the structures almost being superimposable (rmsd of ∼1 Å, [32]). This similarity complicates the prediction of protease activity based on the presence of an intact catalytic triad, since we observed that a catalytic triad can be restored to the *C. difficile* CspA pseudoprotease, yet it does not gain enzymatic activity. This observation raises the possibility that the bioinformatics-based predictions widely used to predict enzyme activity may over-estimate the prevalence of active enzymes. Indeed, a recent review proposed that pseudoenzymes be defined as “the predicted catalytically defective counterparts of enzymes owing to the absence of one or more catatlyic residues[3].” Fortunately, considerable attention is now being directed at improving the curation of pseudoenzymes [48]. Regardless, as more pseudoenzymes are directly studied rather than bioinfomatically predicted, it is likely that additional surprising findings will be made.

Indeed, studying CspA autoprocessing activity in Peptostreptococcaceae family members predicted to encode an active CspA would yield important insights into the evolution of this pseudoprotease and permit assessment of how widely conserved the Csp protease-pseudoprotease signaling system is during clostridial spore germination. These studies may also identify ways to target the *C. difficile* Csp pseudoproteases to prevent spore germination and thus infection, since this signaling system is critical for *C. difficile* to initiate infection [24, 49, 50]. Importantly, pseudoenzymes can be excellent targets for drug discovery because they often have distinct features such as remodeled active sites that can be differentially targeted by small molecules without impacting their active enzyme counterparts [1, 5]. Further elucidating their critical properties and mechanisms of action may therefore represent a promising avenue for therapeutic intervention.

## Acknowledgments

We would like to thank J. Sorg for generously sharing a codon-optimized versions of *cspA, cspB*, and *cspC* and the anti-CspA antibody; N. Minton (U. Nottingham) for providing us with access to the 630Δ*erm*Δ*pyrE* strain and pMTL-YN1C and pMTL-YN3 plasmids for allele-coupled exchange (ACE); and Marcin Dembek for directly providing these materials to us and sharing his specific protocols on ACE with us.

## Declarations of Interest

A.S. has a paid consultancy for BioVector, Inc., a diagnostic start-up.

## Funding Information

Research in this manuscript was funded by Award Number K12 GM133314 to A.E.R., who is a Tufts IRACDA fellow, and T32 GM007310 to E.R.F., and Award Number R01GM108684 to A.S from the National Institutes of General Medical Sciences and R21AI26067 to A.S. from the National Institutes of Allergy and Infectious Disease. A.S. is a Burroughs Wellcome Investigator in the Pathogenesis of Infectious Disease supported by the Burroughs Wellcome Fund. M.L.D was supported in part by the Alpha Omega Alpha Carolyn L. Kuckein Student Research Fellowship. The content is solely the responsibility of the author(s) and does not necessarily reflect the views of the Burroughs Wellcome Fund, or the National Institutes of Health. The funders had no role in study design, data collection and interpretation, or the decision to submit the work for publication.

## Author Contribution Statement

A.S. conceived the hypothesis and supervised the project with help from A.E.R. M.L.D, E.R.F, A.E.R., and A.S designed the experiments. A.S. constructed the single degenerate site mutants for CspC in *C. difficile*, codon-optimized *cspC* mutants carrying single degenerate site mutants, and codon-optimized *cspBA* expression construct. M.L.D. constructed the double degenerate site mutant of CspC, all the CspA degenerate site mutants, and *cspBA* codon-optimized expression constructs encoding degenerate site mutations. E.R.F. cloned the G171R and double degenerate site mutant expression constructs for codon-optimized CspC. M.L.D. performed the phenotypic characterization of *C. difficile* strains (heat-resistance, plate-based and optical density-based germination assays, and western blot analyses) unless otherwise indicated, as well as the *E. coli* protein purification analyses of CspBA. E.R.F. performed the *E. coli* protein purification analyses of CspC. A.S. performed the phenotypic analyses of the *cspBA* prodomain trans-complementation analyses. M.L.D and A.S. wrote the manuscript with help from A.E.R. and E.R.F.

## Figure Legends

**Supplemental figure 1: The CspB prodomain can be supplied *in trans* to reconstitute CspB function and largely complements for loss of CspBA.**

(A) Schematic of wild-type CspBA and a construct encoding the prodomain *in trans* (Q66_TAG_). “Pro” denotes the prodomain. Q66-TAG encodes a variant in which the CspB prodomain is produced *in trans* from the remainder of CspBA through the introduction of a stop codon after codon 66 and a ribosome binding site and start codon before codon 67. (B) Western blot analyses of CspB(A) and CspC in sporulating cells and purified spores from wild type 630Δ*erm*-p, Δ*cspBA*, and Δ*cspBA* complemented with either wild-type *cspBA* or the *cspBA* trans-complementation variant. A-P refers to CspB(A) that has undergone autoprocessing to release the CspB prodomain. Δ*spo0A* (Δ*0A*) was used as a negative control for sporulating cells. SpoIVA was used as a loading control for sporulating cells, while CotA was used as a loading control for purified spores. An anti-CspB antibody was used to detect full-length CspBA in sporulating cells. A non-specific band in the anti-CspB blot is indicated with an asterisk. The germination efficiency of spores from the indicated strains plated on BHIS media containing 0.1% taurocholate is also shown relative to wild type. The mean and standard deviations shown are based on multiple replicates performed on two independent spore purifications. Statistical significance relative to wild type was determined using a one-way ANOVA and Tukey’s test. (C) Germinant sensitivity of Q66_TAG_ spores plated on BHIS containing increasing concentrations of taurocholate. The number of colony forming units (CFUs) produced by germinating spores is shown. The mean and standard deviations shown are based on multiple replicates performed on two independent spore purifications. Statistical significance relative to wild type was determined using a one-way ANOVA and Tukey’s test. **** p < 0.0001, *** p < 0.001, **p < 0.01.

