## Supplementary figures and images for "Differential Effects of “Resurrecting” Csp Pseudoproteases during *Clostridioides difficile* Spore Germination"

### Supplemental Figure 1

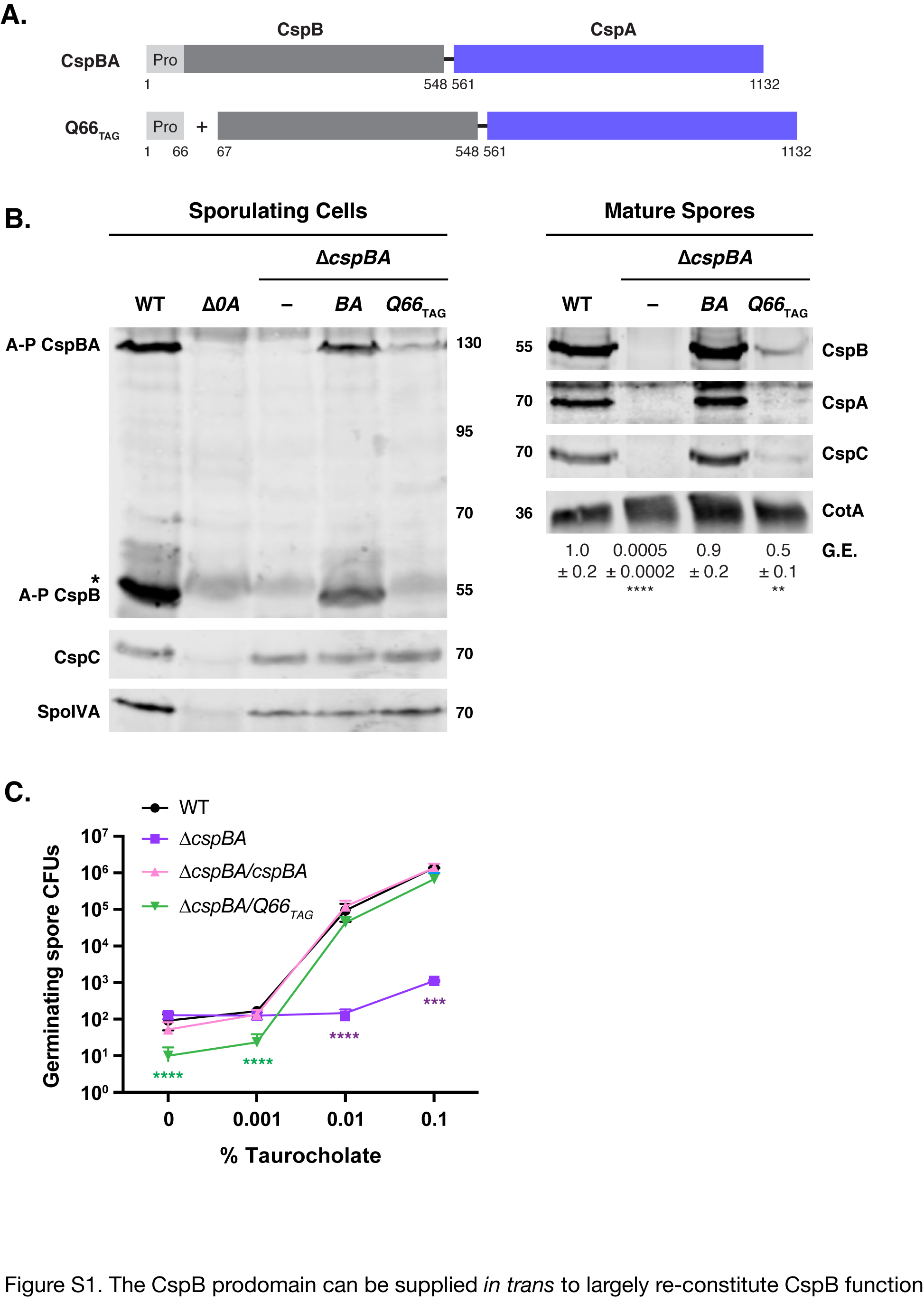
